# Two Chloroflexi classes independently evolved the ability to persist on atmospheric hydrogen and carbon monoxide

**DOI:** 10.1101/457697

**Authors:** Zahra F. Islam, Paul R.F. Cordero, Joanna Feng, Ya-Jou Chen, Sean K. Bay, Thanavit Jirapanjawat, Roslyn M. Gleadow, Carlo R. Carere, Matthew B. Stott, Eleonora Chiri, Chris Greening

**Author notes:** Correspondence can be addressed to: Dr Chris Greening, School of Biological Sciences, Monash University, Clayton, VIC 3800, Australia.

## Abstract

Bacteria within aerated environments often exist within a variety of dormant forms. In these states, bacteria endure adverse environmental conditions such as organic carbon starvation by decreasing metabolic expenditure and using alternative energy sources. In this study, we investigated the energy sources that facilitate the persistence of the environmentally widespread but understudied bacterial phylum Chloroflexi. A transcriptome study revealed that *Thermomicrobium roseum* (class Chloroflexia) extensively remodels its respiratory chain upon entry into stationary phase due to organic carbon limitation. Whereas primary dehydrogenases associated with heterotrophic respiration were downregulated, putative operons encoding enzymes involved in molecular hydrogen (H_2_), carbon monoxide (CO), and sulfur compound oxidation were significantly upregulated. Gas chromatography and microsensor experiments were used to show that *T. roseum* aerobically respires H_2_ and CO at a range of environmentally relevant concentrations to sub-atmospheric levels. Phylogenetic analysis suggests that the enzymes mediating atmospheric H_2_ and CO oxidation, namely group 1h [NiFe]-hydrogenases and type I carbon monoxide dehydrogenases, are widely distributed in Chloroflexi genomes and have been acquired on at least two occasions through separate horizontal gene transfer events. Consistently, we confirmed that the sporulating isolate *Thermogemmatispora* sp. T81 (class Ktedonobacteria) also oxidises atmospheric H_2_ and CO during persistence. This study provides the first axenic culture evidence that atmospheric CO supports bacterial persistence and reports the third phylum to be experimentally shown to mediate the biogeochemically and ecologically important process of atmospheric H_2_ oxidation. This adds to the growing body of evidence that atmospheric trace gases serve as dependable energy sources for the survival of dormant microorganisms.

## Introduction

Bacteria from the phylum Chloroflexi are widespread and abundant in free-living microbial communities [1–4]. One reason for their success is their metabolic diversity; cultured strains from the phylum include heterotrophs, lithotrophs, and phototrophs adapted to both oxic and anoxic environments [5]. Cultured representatives of the phylum are classified into four classes by the genome taxonomy database [6], the primarily aerobic Chloroflexia and Ktedonobacteria and the anaerobic Anaerolineae and Dehalococcoidia [5]. Numerous studies have provided insight into the metabolic strategies anaerobic classes within Chloroflexi use to adapt to oligotrophic niches [7, 8]. Surprisingly little, however, is known about how aerobic heterotrophic bacteria within this phylum colonise oxic environments. Global surveys have reported that Chloroflexi comprise 4.3% of soil bacteria [2] and 3.2% of marine bacteria [3]. However, the most dominant lineages within these ecosystems (notably Ellin6524 and SAR202) have proven recalcitrant to cultivation attempts. Instead, most of our knowledge about the ecophysiological strategies of aerobic heterotrophic Chloroflexi is derived from studies on thermophilic isolates. Various strains from the classes Chloroflexia and Ktedonobacteria have been isolated and characterised from hot springs and geothermal soils [9–13].

Within geothermal environments, Chloroflexi strains must endure temporal and spatial fluxes in the availability of organic carbon compounds. It is currently unknown how members of this phylum stay energised in response to these environmental perturbations. Carbon monoxide (CO) and molecular hydrogen (H_2_) of both geothermal and atmospheric origin are available in such environments and may be particularly important energy sources for sustaining growth and persistence [14–18]. Consistently, genomic and metagenomic studies have revealed that Chloroflexi encode carbon monoxide dehydrogenases [19–21] and hydrogenases [19, 22–24] known to mediate aerobic respiration of these gases. Chloroflexi isolates have been shown to aerobically oxidise CO at a range of environmentally significant concentrations: *Thermomicrobium roseum* can grow chemolithoautotrophically on high concentrations of CO [19] and multiple *Thermogemmatispora* isolates have been shown to oxidise CO, including *T. carboxidovorans* to atmospheric concentrations (0.10 ppmv) [11]. While H_2_ oxidation has yet to be reported in aerobic heterotrophic Chloroflexi, strains of the phylum are known to encode the high-affinity group 1h [NiFe]-hydrogenase [22, 25]. This enzyme class has been shown to support bacterial persistence by mediating oxidation of atmospheric H_2_ [24, 26–34]. To date, atmospheric H_2_ oxidation has only been experimentally confirmed in Actinobacteria [26, 28–30, 32, 35] and two acidobacterial isolates [31, 36].

In this study, we investigated the persistence strategies of thermophilic isolates from two classes of Chloroflexi. We focused primarily on *Thermomicrobium roseum* (class Chloroflexia), a strain originally isolated from Toadstool Spring of Yellowstone National Park, USA [9]. This obligately aerobic bacterium is known to grow heterotrophically on a variety of carbohydrates, organic acids, and proteinaceous substrates, and chemolithoautotrophically on carbon monoxide at high concentrations [9, 12, 19]. Previous genome analyses have shown *T. roseum* encodes a type I carbon monoxide dehydrogenase and a group 1h [NiFe]-hydrogenase [19, 22]. A combination of transcriptome sequencing and targeted activity assays were used to holistically determine the metabolic basis of persistence in this organism, including demonstration of CO and H_2_ oxidation by this strain during carbon-limitation. To generalise these findings, we also investigated *Thermogemmatispora* sp. T81 (class Ktedonobacteria), a cellulolytic thermophilic strain which we previously isolated from geothermal soils in Tikitere, New Zealand [10, 37, 38]. Collectively, our results demonstrate that atmospheric H_2_ and CO serve as important energy sources that support the persistence of this phylum.

## Materials and Methods

### Bacterial strains

*Thermomicrobium roseum* DSM 5159 [9, 12] and *Thermogemmatispora* sp. T81 [10, 37] were imported from the Extremophiles Research Group (GNS Science, Wairakei, New Zealand) culture collection in February 2017. Cultures of both bacterial isolates were routinely maintained in 120 mL serum vials sealed with lab-grade butyl rubber stoppers. Cultures of *T. roseum* cultures contained 30 mL Castenholz media supplemented with 1 g L^−1^ yeast extract and 1 g L^−1^ tryptone, whereas *Thermogemmatispora* sp. T81 cultures were maintained in 30 mL 10% R2A media. For all experiments, both strains were incubated at 60°C at an agitation speed of 150 rpm. Gram staining and 16S rRNA gene amplicon sequencing confirmed both cultures were axenic. Sporulation of *Thermogemmatispora* sp. T81 was verified by light microscopy of Gram-stained cultures.

### Transcriptomics

Transcriptome shotgun sequencing (RNA-Seq) was used to compare gene expression in *T. roseum* cultures under carbon-replete (exponential phase; 10 mL, OD_600_ of 0.3) and carbon-exhausted (stationary phase; 10 mL, OD_600_ of 0.75, 48 hr post OD_max_) conditions. Biological triplicate samples for each condition were harvested by centrifugation (21,000 × *g*, 15 min, 4°C), the supernatants were removed, and the cell pellets resuspended in 1 mL of RNAlater Stabilisation Solution (ThermoFisher Scientific) prior to freezing at −20°C. Extraction and sequencing of RNA was performed by Macrogen Inc., Seoul, Korea. Briefly, RNA was extracted using the RNeasy Plant Mini Kit (Qiagen), libraries were constructed using a TruSeq RNA v2 Sample Prep Kit (Illumina), and rRNA was removed using the Ribo-Zero rRNA Removal Kit (Illumina). The resultant complementary DNA was sequenced on an Illumina HiSeq4000 platform using a paired-end, 100 bp high-throughput protocol. Sequence analysis was performed using the automated cloud-based transcriptomics pipeline, AIR (Sequentia Biotech). Briefly, the steps performed were read trimming, read quality analysis using FastQC [39], and read mapping against the *T. roseum* reference genome (NCBI ID: NC_011959.1 [19]) using an intrinsic platform read aligner with default parameters. Aligned reads were checked for quality prior to data normalisation using the trimmed mean of M-values method [40] and the ‘normalizaData’ command within the R package HTSFilter [41]. A principle component analysis was then performed on the normalised data prior to statistical analysis using edgeR [42] to obtain differential gene expression counts.

### Gas chromatography

Gas chromatography was used to determine whether the two Chloroflexi strains were capable of oxidising atmospheric levels of CO and H_2_. Briefly, sealed serum vials containing stationary-phase cultures of *T. roseum* (72 hr post OD_max_) and sporulating cultures of *Thermogemmatispora* sp. T81 (294 hr post-inoculation) were opened, aerated, and resealed. They were then amended with H_2_ (*via* 1% v/v H_2_ in N_2_ gas cylinder, 99.999% pure) or CO (*via* 1% v/v CO in N_2_ gas cylinder, 99.999% pure) to achieve headspace concentrations of ~14 ppmv. Cultures were agitated (150 rpm) for the duration of the incubation period to enhance H_2_ and CO transfer to the cultures and maintain an aerobic environment. Headspace samples (1 mL) were routinely collected using a gas-tight syringe to measure H_2_ and CO. Concomitantly, headspace gas concentrations in heat-killed negative controls (30 mL) were measured to confirm that observed rates of gas consumption occurred due to a biotic process. Gas concentrations in samples were measured by gas chromatography using a pulsed discharge helium ionization detector [43, 44]. This customized trace gas analyser (model TGA-6791-W-4U-2, Valco Instruments Company Inc.) is designed to analyse a suite of atmospheric gases across six orders of magnitude in concentrations. Briefly, the system is configured to use two valves as injectors/backflushers, and two valves to front flush or heart cut from the precolumns (Mole Sieve 5A, set at 140°C). Gases are then separated on the main columns (5’ X 1/8” HayseSep Db, set at 55°C). The fifth valve is used as a sample loop selector to accommodate a larger range of gas concentrations. Concentrations of H_2_ and CO in each sample were regularly calibrated against ultra-pure H_2_ and CO gas standards of known concentrations. With the standards used, the limit of detection was 42 ppbv H_2_ and 9 ppbv CO.

### Kinetic analysis

Apparent kinetic parameters of H_2_ and CO oxidation were estimated in stationary-phase *T. roseum* cultures. Briefly, cultures were incubated with 100 or 1000 ppmv H_2_ and 200 or 1500 ppmv CO, and headspace gas samples were measured at various time intervals (0, 2, 4 and 8 hr from substrate addition) by gas chromatography. Reaction velocity relative to the gas concentration was measured at each timepoint and plotted on a Michaelis-Menten curve. Curves of best fit, *V*_max_ app values, and *K*_m_ app values were calculated in GraphPad Prism (version 7.01) using non-linear regression models (enzyme kinetics – substrate vs. velocity, Michaelis-Menten, least squares fit).

### Activity staining

Hydrogenase and carbon monoxide dehydrogenase activity was stained using whole-cell lysates of stationary-phase cultures of *T. roseum*. Five hundred mL of culture was harvested by centrifugation (10,000 *× g*, 10 min, 4°C), washed in phosphate-buffered saline solution (PBS; 137 mM NaCl, 2.7 mM KCl, 10 mM Na_2_HPO_4_ and 2 mM KH_2_PO_4_, pH 7.4), and resuspended in 16 mL lysis buffer (50 mM MOPS, pH 7.5, 1 mM PMSF, 15 mM MgCl_2_, 5 mg mL^−1^ lysozyme, 1 mg DNase). The resultant suspension was then lysed by passage through a Constant Systems cell disruptor (40,000 psi, four times), with unbroken cells removed by centrifugation (10,000 *× g*, 20 min, 4°C). Protein concentration was calculated using the bicinchoninic acid assay [45] against bovine serum albumin standards. Next, 20 μg protein was loaded onto native 7.5% (w/v) Bis-Tris polyacrylamide gels prepared as described elsewhere [46] and run alongside a protein standard (NativeMark Unstained Protein Standard, Thermo Fisher Scientific) at 25 mA for 3 hr. For total protein staining, gels were incubated in AcquaStain Protein Gel Stain (Bulldog Bio) at 4°C for 3 hr. For hydrogenase staining [47], gels were incubated in 50 mM potassium phosphate buffer supplemented with 500 μM nitroblue tetrazolium chloride (NBT) in an anaerobic chamber (5% H_2_, 10% CO_2_, 85% N_2_ v/v) at 60°C for 4 hr. For carbon monoxide dehydrogenase staining [48], gels were incubated in 50 mM Tris-HCl buffer containing 50 μM NBT and 100 μM phenazine methosulfate in an anaerobic chamber (100% CO v/v atmosphere) maintained at 60°C for 4 hr.

### Electrode measurements

For *T. roseum* cultures, rates of H_2_ oxidation with and without treatment of respiratory chain uncouplers were measured amperometrically, following previously established protocols [49, 50]. Prior to the start of measurement, a Unisense H_2_ microsensor electrode was polarised at +800 mV for 1 hr using a Unisense multimeter and calibrated with standards of known H_2_ concentration. Gas-saturated PBS was prepared by bubbling the solution with 100% (v/v) of either H_2_ or O_2_ for 5 minutes. For untreated cells, 1.1 mL microrespiration assay chambers were sequentially amended with stationary-phase *T. roseum* cell suspensions (OD_600_ = 1; 0.9 mL), H_2_-saturated PBS (0.1 mL), and O_2_-saturated PBS (0.1 mL) stirred at 250 rpm, 37°C. Following measurements of untreated cells, the assay mixtures were treated with 100 μM carbonyl cyanide *m*-chlorophenyl hydrazine (CCCP), 10 μM nigericin, or 10 μM valinomycin. Changes in H_2_ concentrations were recorded using Unisense Logger Software (agitation speed of 250 rpm, 37°C). Upon observing a linear change in H_2_ concentration, initial rates of consumption were calculated over a period of 20 s and normalised against total protein concentration.

### Phylogenetic analyses

Phylogenetic trees were constructed to investigate the evolutionary history and distribution of uptake hydrogenases and carbon monoxide dehydrogenases within the Chloroflexi phylum. Specifically, the catalytic subunits of [NiFe]-hydrogenases (HhyL and homologues) and type I carbon monoxide dehydrogenases (CoxL) were retrieved from Chloroflexi genomes and metagenome-assembled genomes (MAGs) in the NCBI RefSeq database *via* protein BLAST [51] in October 2018. The amino acid sequences were aligned with reference sequences [21, 22] using ClustalW in MEGA7 [52].Evolutionary relationships were visualised by constructing a maximum-likelihood phylogenetic tree in MEGA7 [52]. Specifically, initial trees for the heuristic search were obtained automatically by applying Neighbour-Join and BioNJ algorithms to a matrix of pairwise distances estimated using a JTT model, and then selecting the topology with superior log likelihood value. Gaps were treated with partial deletion and trees were bootstrapped to 100 replicates.

## Results and Discussion

### *Thermomicrobium roseum* upregulates hydrogenase and carbon monoxide dehydrogenase expression during a coordinated response to organic carbon starvation

We compared the transcriptomes of triplicate *T. roseum* cultures under carbon-replete (exponential growing) and carbon-limited (stationary phase) conditions. 401 genes were significantly upregulated and 539 genes were significantly downregulated by at least two-fold (*p* < 10^−6^) in response to carbon-limitation (**Figure 1a**; **Table S1**). Three major trends were observed with respect to energy acquisition and utilisation. Firstly, genes associated with energetically expensive processes were downregulated, including those encoding ribosomal proteins, cytochrome *c* and menaquinone biosynthesis enzymes, and the megaplasmid-encoded chemotactic and flagellar apparatus (**Table S1**). Secondly, there was evidence of mobilisation of internal carbon stores, including an acetoin dehydrogenase complex and an electron transfer flavoprotein complex (ETF). Thirdly, primary respiratory dehydrogenases involved in heterotrophic growth (i.e. type I and II NADH dehydrogenases) were downregulated in favour of those associated with lithotrophic metabolism (**Figure 1a**; **Table S1**). In both conditions, the terminal oxidases that mediate aerobic respiration were highly expressed and there was no evidence of the use of other electron acceptors; the cytochrome *aa*3 oxidase was expressed in both phases and the alternative cytochrome *bo*3 oxidase was upregulated during stationary-phase. In contrast, the F_1_F_o_-ATPase (ATP synthase) was downregulated, a finding consistent with an expected decrease in the availability of respiratory electron donors during carbon-limitation (**Table S1**).

**Figure 1.**
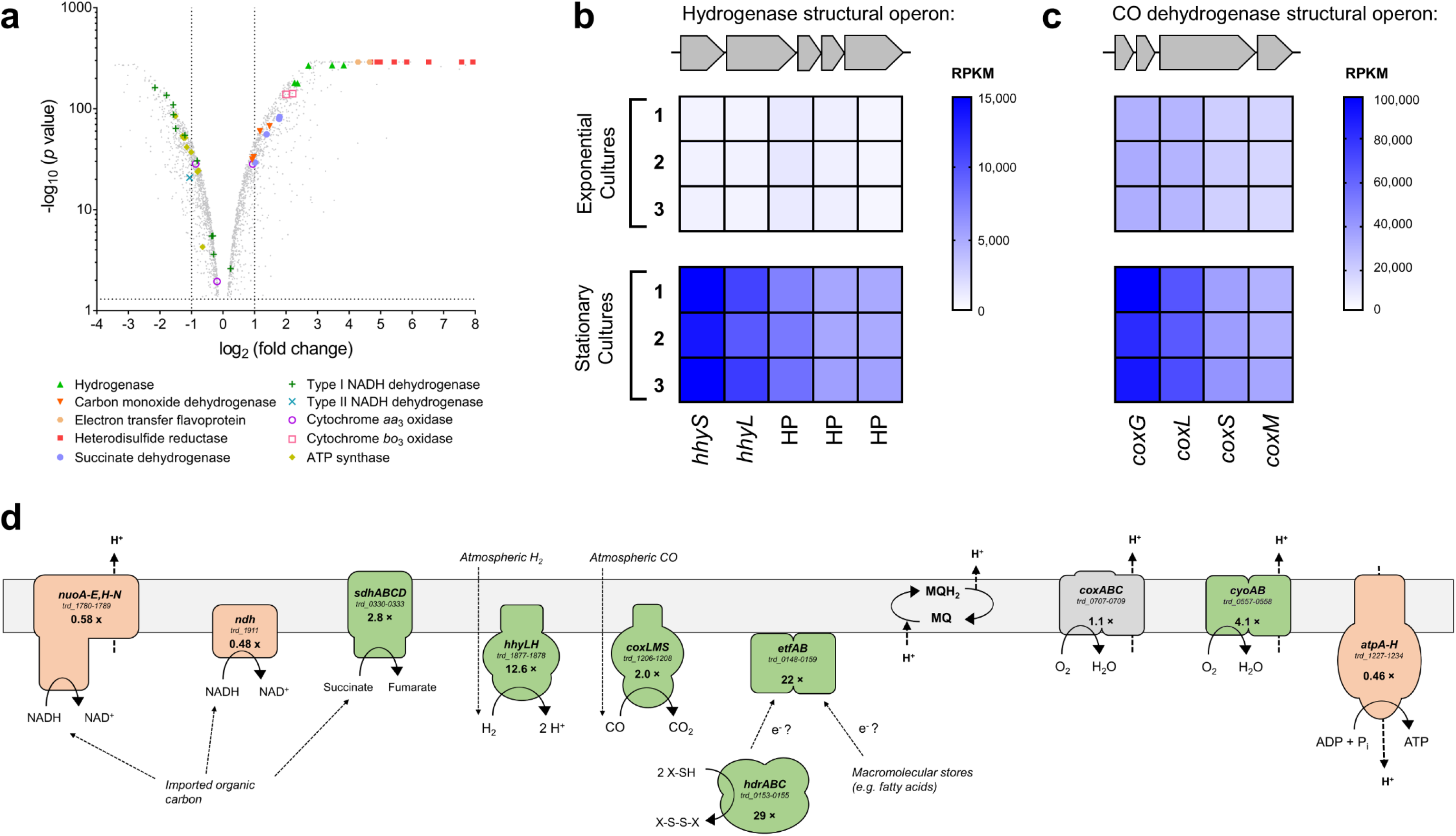
Differential gene expression of carbon-replete (exponential phase) and carbon-limited (stationary phase) cultures of *Thermomicrobium roseum.* (**a**) Volcano plot showing relative expression change of genes following carbon-limitation. The fold-change shows the ratio of normalised transcript abundance of three stationary phase cultures divided by three exponential phase cultures (biological replicates). Each gene is represented by a grey dot and respiratory genes are highlighted as per the legend. (**b & c**) Heat maps of normalised abundance of the putative operons encoding the structural subunits of the group 1h [NiFe]-hydrogenase (*hhySL*; **a**) and type I carbon monoxide dehydrogenase (*coxLSM*; **b**). The read counts per kilobase million (RPKM) are shown for three exponentially growing and three stationary phase biological replicates. (**d**) Differential regulation of the respiratory complexes mediating aerobic respiration of organic and inorganic compounds. Complexes that are differentially shaded depending on whether they are significantly upregulated (green), downregulated (orange), or unchanged (grey) in carbon-limited compared to carbon-replete cultures. Gene names, loci numbers, and average fold changes in transcriptome abundance are shown for each complex. Note that the physiological role of the highly upregulated *hdr* and *etf* complexes is yet to be experimentally validated.

*Thermomicrobium roseum* upregulates genes associated with H_2_ and CO metabolism under carbon-limiting conditions. The genes encoding the structural subunits of a group 1h [NiFe]-hydrogenase (*hhySL*; trd_1878-1877) [22, 23, 53], which is known to mediate atmospheric H_2_ oxidation [29, 31, 47], were upregulated by an average of 12.6-fold (**Figure 1b**). Also upregulated were the conserved hypothetical proteins *hhaABC* (trd_1876-1874; 5.5-fold) [25], encoded on the same putative operon as the structural subunits, as well as a separate putative operon of maturation factors (trd_1873-1863; 3.1-fold) (**Figure S1; Table S1**). The structural (trd_1206-1208) and maturation (trd_1209-1215) subunits encoding a type I carbon monoxide dehydrogenase were upregulated by an average of twofold (**Figure 1c** **& S1**) in response to carbon limitation. Consistent with previous reports of chemolithoautotrophic growth at high concentrations of CO [19], carbon monoxide dehydrogenase genes were highly expressed in both exponential and stationary phase cultures. (**Figure 1c**; **Table S1**). This suggests that *T. roseum* uses CO to supplement available organic carbon during growth (mixotrophy) and persistence.

Overall, the greatest differential in gene expression involved a 19-gene cluster (trd_0160-0142) putatively involved with the oxidation of sulfur compounds. The cluster encodes a soluble heterodisulfide reductase (*hdrABC*), an electron transfer flavoprotein complex (*etfAB*), three sulfur carriers (*dsrE1, dsrE2, tusA*), three lipoate-binding proteins (*lbpA*), and various hypothetical proteins, which are upregulated by an average of 45-fold during persistence. We predict that this system serves to mobilise sulfur atoms from inorganic or organic sulfur-containing compounds and conjugate them to sulfur carriers. In turn, we hypothesise that the Hdr complex may oxidise the resultant thiol groups to disulfides and transfer the liberated electrons to the aerobic respiratory chain-linked ETF complex. In support of this premise, the upregulated gene cluster encodes components homologous to a system recently shown to mediate the oxidation of diverse sulfur compounds in *Hyphomicrobium denitrificans* [54, 55] and the Hdr complex is most closely related to those of sulfur-oxidising *Sulfobacillus, Hyphomicrobium,* and *Acidithiobacillus* strains. It seems plausible that *T. roseum* would benefit from a survival advantage if able to harness reduced sulfur compounds available in geothermal springs. However, further work is needed to verify the activity, substrates, and physiological role of this system.

Collectively, these findings show that *T. roseum* is more metabolically flexible than previously thought. **Figure 1d** illustrates the predicted remodelling of the respiratory chain that occurs during the transition from carbon-replete to carbon-limited conditions. The upregulation of enzymes involved in harnessing inorganic compounds, in conjunction with the downregulation of gene clusters involved in organic carbon oxidation, suggests that *T. roseum* has evolved mechanisms to maintain aerobic respiration despite organic carbon fluctuations and deprivation within its environment.

### *T. roseum* aerobically oxidises H_2_ and CO at a wide range of concentrations, including sub-atmospheric levels, during persistence

The high expression levels for genes encoding the group 1h [NiFe]-hydrogenase and type I carbon monoxide dehydrogenase suggested that *T. roseum* may support persistence by oxidising atmospheric H_2_ and CO. To test this, we incubated carbon-limited cultures of *T. roseum* in an ambient air headspace supplemented with ~14 ppmv of either H_2_ or CO and monitored their consumption using gas chromatography. In agreement with our hypothesis, cultures aerobically oxidised both gases in a firstorder kinetic process and reached mixing ratios (103 ppbv H_2_, 22 ppbv CO) five times below atmospheric levels within 71 hours (**Figure 2a & b**). This constitutes the first observation of both aerobic H_2_ respiration and atmospheric H_2_ oxidation within the phylum Chloroflexi.

**Figure 2.**
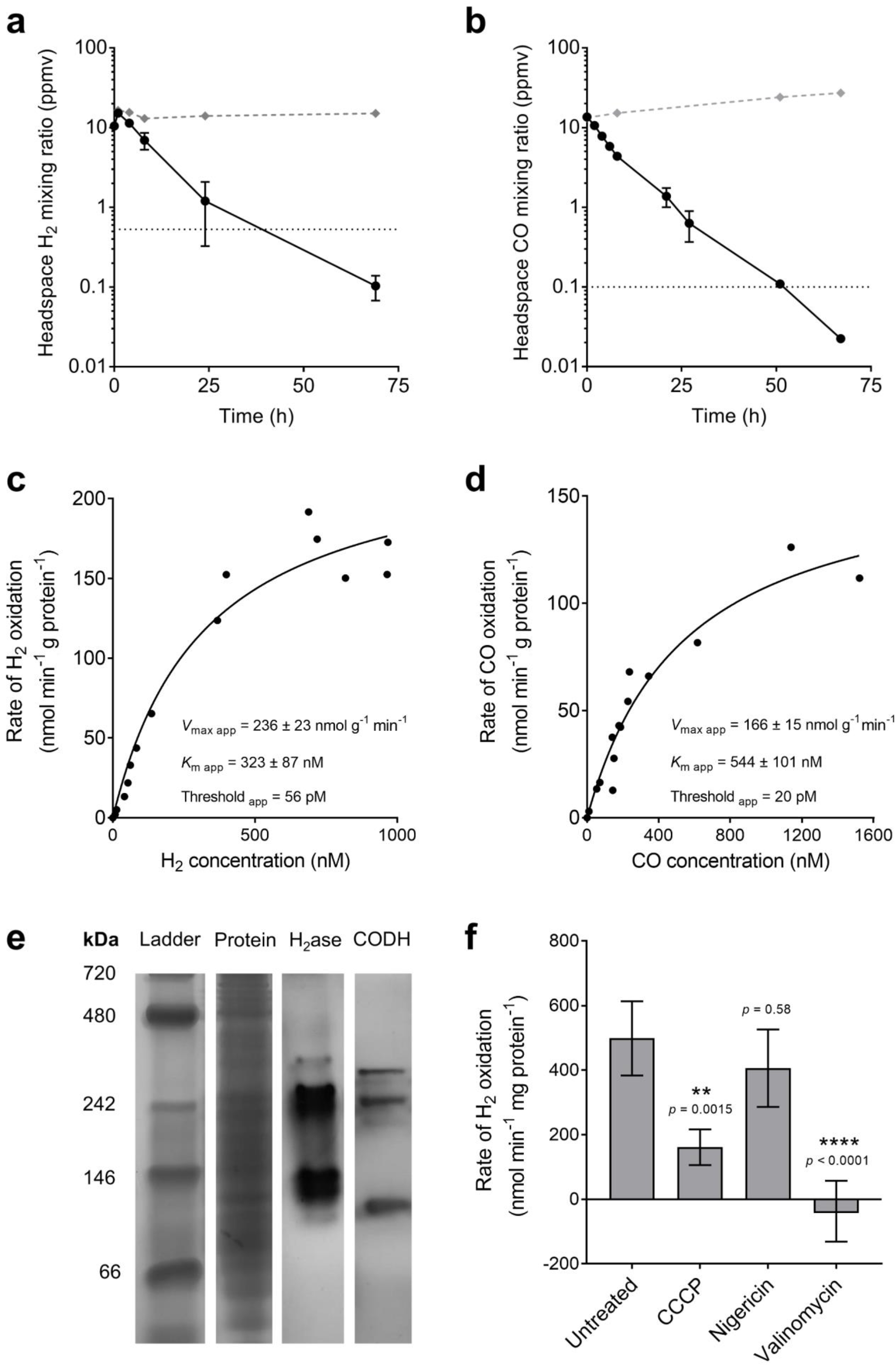
Hydrogenase and carbon monoxide dehydrogenase activity of *Thermomicrobium roseum* cultures during organic carbon limitation. (**a & b**) Oxidation of molecular hydrogen (H_2_; **a**) and carbon monoxide (CO; **b**) to sub-atmospheric levels by *T. roseum* cultures. Error bars show standard deviations of three biological replicates, with heat-killed cells monitored as a negative control. Mixing ratios of H_2_ and CO are displayed on a logarithmic scale and dotted lines show the average atmospheric mixing ratios of H_2_ (0.53 ppmv) and CO (0.10 ppmv). (**c & d**) Apparent kinetics of H_2_ (**c**) and CO (**d**) oxidation by *T. roseum* whole cells. Curves of best fit and kinetic parameters were calculated based on a Michaelis-Menten kinetics model. (**e**) Zymographic observation of hydrogenase and carbon monoxide dehydrogenase activity in *T. roseum* whole-cell lysates. The first two lanes show protein ladder and whole protein stained with Coomassie Blue. The other lanes shown hydrogenase and carbon monoxide dehydrogenase activity stained with the artificial electron acceptor nitroblue tetrazolium in a H_2_-rich and CO-rich atmosphere respectively. (**f**) Amperometric measurements of hydrogenase activity in *T. roseum* whole cells. The rate of H_2_ oxidation was measured with a hydrogen electrode before and after treatment with the respiratory uncouplers and ionophores carbonyl cyanide *m*-chlorophenyl hydrazine (CCCP), nigericin, and valinomycin.

Whole-cell kinetic measurements revealed that *T. roseum* efficiently oxidises H_2_ and CO across a wide range of concentrations through hydrogenase and carbon monoxide dehydrogenase activity. The enzymes display a moderate apparent velocity (*V*_max_ app of 236 nmol H_2_ and 166 nmol CO g^−1^ of protein min^−1^) and moderate apparent affinity (*K*_m_ app of 323 nM H_2_ and 544 nM CO) for these substrates (**Figure 2c & d**). With respect to carbon monoxide dehydrogenase, these observations are consistent with the organism being able to utilise CO at elevated concentrations for growth [19] and atmospheric concentrations for persistence. The apparent kinetic parameters of the group 1h [NiFe]-hydrogenase are more similar to those recently described for the verrucomicrobial methanotroph *Methylacidiphilum fumariolicum* (*K*_m_ = 600 nM) [56] than to the high-affinity, low-activity hydrogenases of previously described atmospheric H_2_ scavengers (*K*_m_ < 50 nM) [26, 29, 31]. Altogether, these findings suggest that *T. roseum* can take advantage of the elevated H_2_ and CO concentrations when available through geothermal activity and subsist on atmospheric concentrations of these gases otherwise.

Consistent with observed whole-cell activities, cell-lysates run on native polyacrylamide gels strongly stained for hydrogenase and carbon monoxide dehydrogenase activity (**Figure 2e**). The triplet banding pattern observed for the hydrogenase is similar to that shown to be mediated by the group 1h [NiFe]-hydrogenases in *M. smegmatis* [26]. We next verified that the hydrogenase was coupled to the respiratory chain by measuring H_2_ oxidation using a H_2_ electrode under aerobic conditions. Untreated cells oxidised H_2_ at a rapid rate. This activity decreased by 2.5-fold upon addition of the respiratory uncoupler CCCP and ceased upon addition of the ionophore valinomycin (**Figure 2f**).

Findings from the transcriptome analysis and activity studies therefore suggest that *T. roseum* persists through oxidation of atmospheric H_2_ and CO. We propose that the group 1h [NiFe]-hydrogenase and type I carbon monoxide dehydrogenase directly use electrons derived from atmospheric H_2_ and CO to support aerobic respiration (**Figure 1d**). Due to the genetic intractability of Chloroflexi and the lack of specific hydrogenase or carbon monoxide dehydrogenase inhibitors, we were unable to determine the necessity of either H_2_ or CO oxidation for prolonged survival for this organism. However, previous studies have demonstrated that genetic deletion of the group 1h [NiFe]-hydrogenase reduces longevity of *M. smegmatis* cells [27, 28, 57] and *Streptomyces avermitilis* exospores [29, 30].

### Scavenging of atmospheric gases is a general persistence strategy within the aerobic heterotrophic Chloroflexi

Having demonstrated that *T. roseum* oxidises atmospheric trace gases during persistence, we subsequently investigated whether this is a common strategy in employed by the Chloroflexi. We first analysed the respiratory capabilities of *Thermogemmatispora* sp. T81, a heterotrophic cellulolytic and spolulating thermophile which we previously isolated from geothermal soils from Tikitere, New Zealand [10, 38]. Analysis of the organism’s genome (Assembly ID: GCA_003268475.1) indicated that it encodes core respiratory chain components similar to *T. roseum*, including primary dehydrogenases (*nuo, ndh, sdh*), terminal oxidases (*cox, cyo*), and ATP synthase (*atp*). The genome also encodes putative operons for the structural subunits of a group 1h [NiFe]-hydrogenase, the maturation factors of this hydrogenase, and structural subunits of a type I carbon monoxide dehydrogenase (**Figure S2**). However, homologues of the putative heterodisulfide reductase and ETF complexes encoded by *T. roseum* are absent from the *Thermogemmatispora* sp. T81 genome.

We verified that sporulating cultures of *Thermogemmatispora* sp. T81 actively consume H_2_ and CO. The organism slowly oxidised available H_2_ and CO in the headspace to sub-atmospheric levels (120 ppbv H_2_, 70 ppbv CO) over ~320 hr (**Figure 3a & b**). Although this strain has previously been shown to oxidise carbon monoxide [11], this is the first observation that it can do so to sub-atmospheric concentrations and during persistence. These results suggest that, despite their distinct evolutionary histories and ecological niches, *Thermogemmatispora* sp. T81 and *T. roseum* have both evolved similar metabolic strategies to survive organic carbon limitation.

**Figure 3.**
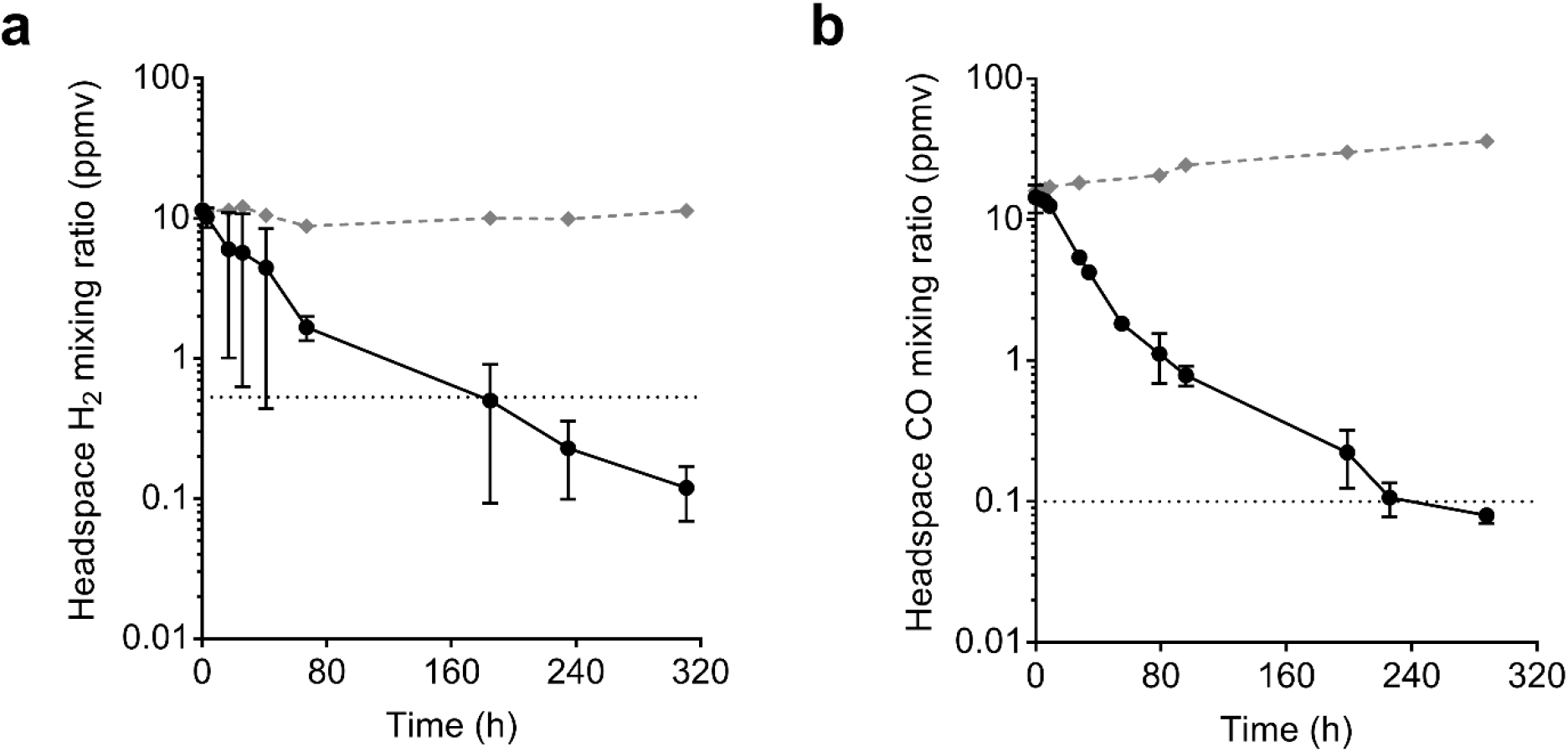
Hydrogenase and carbon monoxide dehydrogenase activity of *Thermogemmatispora* sp. T81. Oxidation of molecular hydrogen (H_2_; **a**) and carbon monoxide (CO; **b**) to sub-atmospheric levels by *Thermogemmatispora* sp. T81 cultures. Error bars show standard deviations of three biological replicates, with heat-killed cells monitored as a negative control. Mixing ratios of H_2_ and CO are displayed on a logarithmic scale and dotted lines show the average atmospheric mixing ratios of H_2_ (0.53 ppmv) and CO (0.10 ppmv).

Analysis of the distribution of hydrogenases and carbon monoxide dehydrogenases within publicly available reference genomes showed that genetic capacity for trace gas scavenging is a common trait among aerobic Chloroflexi. Specifically, group 1 h [NiFe]-hydrogenases and type I carbon monoxide hydrogenases sequences were encoded in three of the four reference genomes within the Thermomicrobiales (class Chloroflexia) and four of the five reference genomes within the Ktedonobacteriales (class Ktedonobacteria) (**Figure 4a & b**). The latter includes the genomes of the heterotrophic soil bacterium *Ktedonobacter racemifer* [58] and the nitrite-oxidising bioreactor isolate *Nitrolancea hollandica* [59]. In addition, seven strains within the photosynthetic order Chloroflexales encoded group 1f and/or group 2a [NiFe]-hydrogenases (**Figure S3**). These hydrogenase classes have been shown to mediate aerobic H_2_ oxidation in a range of bacteria, including to sub-atmospheric concentrations in *Acidobacterium ailaaui* and *M. smegmatis* respectively [36, 47]. Moreover, a metatranscriptome study showed homologs of the group 1f [NiFe]-hydrogenase of *Roseiflexus* species are highly expressed in geothermal microbial mats at night [60]. Hence, it is likely that the traits of aerobic H_2_ respiration and possibly atmospheric H_2_ oxidation extends to the photosynthetic strains of this phylum. A range of metagenome-assembled genomes, including from the abundant candidate class Ellin6549 [2, 24], also encoded genes for aerobic H_2_ and CO oxidation (**Figure S3 & S4**). Consistent with previous reports, Dehalococcoidia encode group 1a [NiFe]-hydrogenases known to facilitate dehalorespiration [61–63].

**Figure 4.**
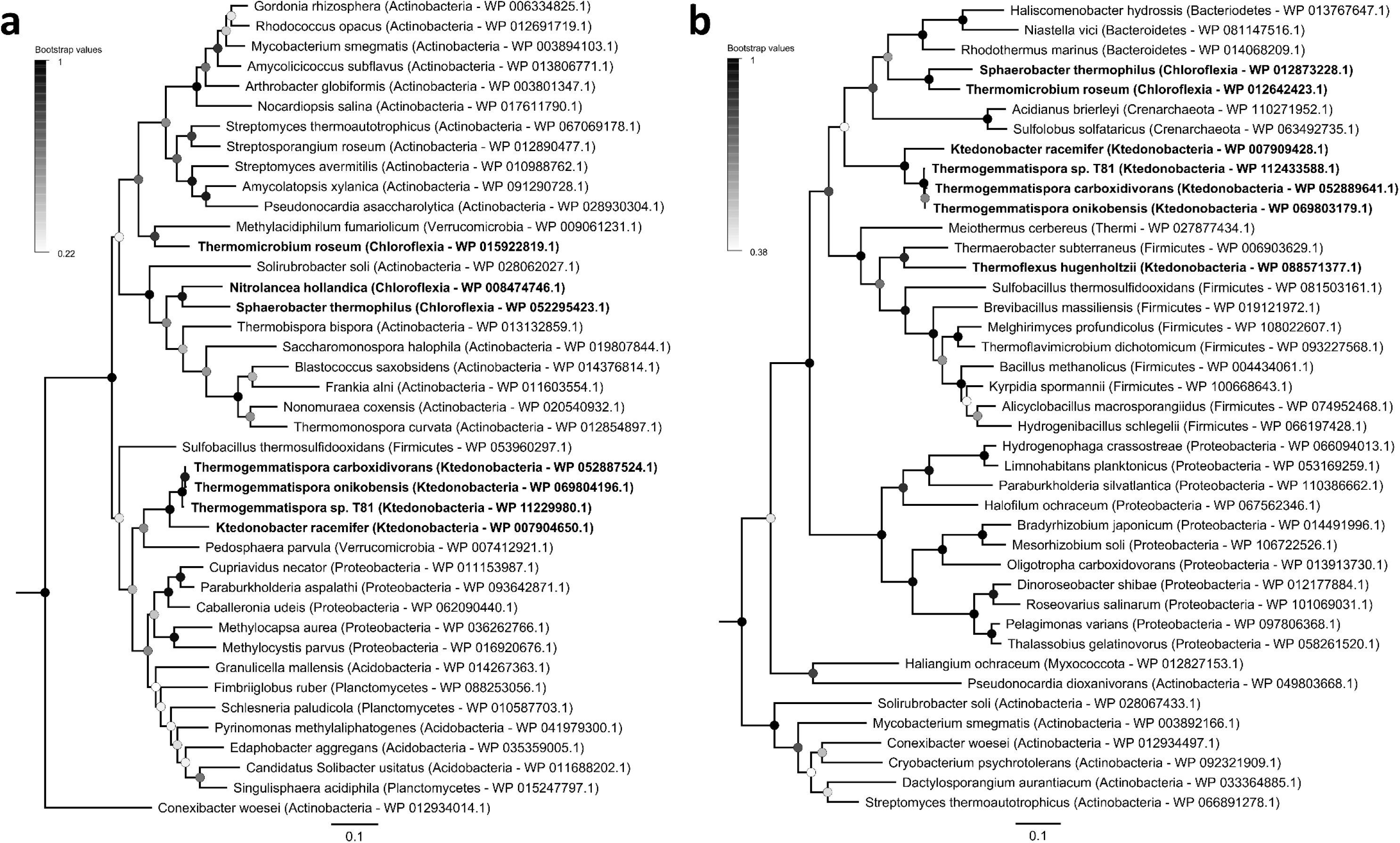
Evolutionary history of the group 1h [NiFe]-hydrogenase and type I carbon monoxide dehydrogenase. Phylogenetic trees showing the distribution and evolutionary history of the catalytic (large) subunits of the group 1h [NiFe]-hydrogenase (*hhyL*; **a**) and type I carbon monoxide dehydrogenase (*coxL*; **b**) in the phylum Chloroflexi. Chloroflexi sequences (labelled by class) are shown in bold against reference sequences (labelled by phylum). Trees were constructed using amino acid sequences through the maximum-likelihood method (gaps treated with partial deletion) and were bootstrapped with 100 replicates. The trees were respectively rooted with group 1g [NiFe]-hydrogenase sequences (WP_011761956.1, WP_048100713.1) and type II carbon monoxide dehydrogenase sequences (WP_011388721.1, WP_012893108.1). The distribution of other respiratory uptake hydrogenases within genomes and metagenome-assembled genomes (MAGs) in the phylum Chloroflexi is shown in **Figure S3.** The distribution of type I carbon monoxide dehydrogenases within metagenome-assembled genomes (MAGs) in the phylum Chloroflexi is shown in **Figure S4.**

Our analyses suggest that the capacity for atmospheric H_2_ and CO oxidation has evolved on at least two occasions within the Chloroflexi. Phylogenetic trees show that the group 1h [NiFe]-hydrogenases from Chloroflexia and Ktedonobacteria are divergent and fall into two distinct, robustly supported branches (**Figure 4a**). It is therefore likely that Chloroflexia and Ktedonobacteria independently acquired these enzymes *via* horizontal gene transfer events from other Terrabacteria, rather than vertically inheriting them from a common ancestor. Phylogenetic analysis also suggests that the type I carbon monoxide dehydrogenase has been acquired on two or three occasions in this phylum (**Figure 4b**). In line with being independently acquired, the putative operons encoding the hydrogenase and carbon monoxide dehydrogenase in *T. roseum* (**Figure S1**) and *Thermogemmatispora* sp. T81 (**Figure S2**) are distinctly organized. For example, the structural and accessory factors of carbon monoxide dehydrogenase are encoded in a single putative operon in *Thermogemmatispora* sp. T81 (*coxMSLIG*), but are separated into a structural operon (*coxGSLM*) and accessory operon (including *coxG* and *coxE*) in *T. roseum*. These findings agree with previous inferences of horizontal dissemination of *hhyL* and *coxL* genes [21, 22, 25] and suggest there is strong selective pressure for the acquisition of metabolic enzymes that support persistence.

### Ecological significance of metabolic flexibility and trace gas oxidation in Chloroflexi

Aerobic heterotrophic bacteria from the phylum Chloroflexi are more metabolically versatile than previously thought. The transcriptome analyses clearly shows that *T. roseum* regulates its metabolism in response to organic carbon limitation, enabling persistence on a combination of exogenous inorganic compounds and likely endogenous carbon reserves. In support of this, gas chromatography measurements showed that the bacterium efficiently oxidises H_2_ and CO down to sub-atmospheric concentrations during persistence through an aerobic respiratory process. We made similar findings for the ktedonobacterial isolate *Thermogemmatispora* sp. T81, suggesting that trace gas scavenging might be a common persistence strategy employed by aerobic Chloroflexi. Phylogenetic analyses suggest that *T. roseum* and *Thermogemmatispora* sp. T81 horizontally acquired the capacity to oxidise atmospheric H_2_ and CO *via* separate events. The apparent convergence in persistence strategies is notable given the distinct evolutionary histories, persistence morphologies (i.e. sporulation in T81), and ecological niches of these bacteria. Resource generalism is therefore likely to be a common ecological strategy for the survival of Chloroflexi in environments where organic carbon may be periodically scarce.

More broadly, these findings provides pure culture support for the hypothesis that atmospheric carbon monoxide serves as an energy source for persistence [24]. A range of heterotrophic bacteria have previously been inferred to be capable of oxidising atmospheric CO, including Proteobacteria [64, 65], Actinobacteria [66], and a *Thermogemmatispora* strain [11]; however, the exact physiological role of this process has remained unresolved. We show that the expression and activity of carbon monoxide dehydrogenase is linked to persistence, and provide evidence that atmospheric CO serves as an electron donor for the aerobic respiratory chain in this condition. Indeed, as with atmospheric H_2_, atmospheric CO is likely to be a dependable energy source for microbial survival given its ubiquity, diffusibility, and energy density. However, in contrast to atmospheric H_2_ [27, 30], it remains to be validated through genetic and biochemical studies that atmospheric CO oxidation can enhance survival of bacteria during persistence. It should also be emphasised that the wide kinetic range of the *T. roseum* carbon monoxide dehydrogenase likely enables this isolate to both persist on atmospheric CO and to grow on CO chemolithoautotrophically [19] (and possibly mixotrophically) within specific microenvironments where carbon monoxide is available at elevated concentrations through geothermal activity.

Finally, this study establishes Chloroflexi as the third phylum experimentally shown to scavenge atmospheric H_2_, following the Actinobacteria [29, 32, 35, 47, 53] and Acidobacteria [31, 36]. The findings made here are similar to those previously reported for the actinobacterium *Mycobacterium smegmatis* [47, 57] and acidobacterium *Pyrinomonas methylaliphatogenes* [31], both of which also shift from heterotrophic respiration to atmospheric H_2_ oxidation in response to energy limitation, including through expressing group 1h [NiFe]-hydrogenases. Given at least four other cultured phyla (**Figure 4a**) and two candidate phyla [24] also encode group 1h [NiFe]-hydrogenases, it seems increasingly likely that atmospheric H_2_ serves as a general energy source for aerobic heterotrophic bacteria. This observation is also potentially biogeochemically significant, given aerobic soil bacteria are known to be the main sink in the global biogeochemical hydrogen cycle [67]. Further work, however, is needed to test these whether these principles extend to the still enigmatic Chloroflexi species inhabiting mesophilic soil environments.

## Footnotes

### Author contributions

C.G. conceived this study, designed experiments, and analysed data. Z.F.I., P.R.F.C., and J.F. performed experiments and analysed data. C.G., R.M.G., and Z.F.I. supervised students. Y.J.-C., S.K.B., T.J., C.R.C., M.B.S., and E.C. contributed to methods development and data analysis. C.G., Z.F.I., C.R.C., M.B.S., and E.C. wrote and edited the paper.

## Acknowledgements

This work was supported by an ARC DECRA Fellowship (DE170100310; awarded to C.G.), an ARC Discovery Project (DP180101762; awarded to C.G.), a Marsden Grant (16-GNS-035; awarded to C.R.C. and C.G.), an SNF Early Mobility Postdoctoral Fellowship (P2EZP3_178421; awarded to E.C.), an ARC Research Training Program Stipend (awarded to Z.F.I.) and Monash University Doctoral Scholarships (awarded to P.R.F.C., Y.-J.C., and S.B.). We thank Prof Craig White and Prof Gary King for their helpful comments.

The authors declare no conflict of interest.

## References

1. Thompson LR, Sanders JG, McDonald D, Amir A, Ladau J, Locey KJ, et al. A communal catalogue reveals Earth’s multiscale microbial diversity. Nature 2017; 551: 457–463.

2. Delgado-Baquerizo M, Oliverio AM, Brewer TE, Benavent-González A, Eldridge DJ, Bardgett RD, et al. A global atlas of the dominant bacteria found in soil. Science 2018; 359: 320–325.

3. Sunagawa S, Coelho LP, Chaffron S, Kultima JR, Labadie K, Salazar G, et al. Structure and function of the global ocean microbiome. Science 2015; 348: 1261359.

4. Mehrshad M, Salcher MM, Okazaki Y, Nakano S, Šimek K, Andrei A-S, et al. Hidden in plain sight—highly abundant and diverse planktonic freshwater Chloroflexi. Microbiome 2018; 6: 176.

5. Whitman WB. Bergey’s manual of systematics of Archaea and Bacteria. 2015. Wiley Online Library.

6. Parks DH, Chuvochina M, Waite DW, Rinke C, Skarshewski A, Chaumeil P-A, et al. A standardized bacterial taxonomy based on genome phylogeny substantially revises the tree of life. Nat Biotechnol 2018; 36: 996–1004.

7. Hug LA, Castelle CJ, Wrighton KC, Thomas BC, Sharon I, Frischkorn KR, et al. Community genomic analyses constrain the distribution of metabolic traits across the Chloroflexi phylum and indicate roles in sediment carbon cycling. Microbiome 2013; 1: 22.

8. Dong X, Greening C, Rattray JE, Chakraborty A, Chuvochina M, Mayumi D, et al. Potential for microbial anaerobic hydrocarbon degradation in naturally petroleum-associated deep-sea sediments. bioRxiv 2018; 400804.

9. Jackson TJ, Ramaley RF, Meinschein WG. Thermomicrobium, a new genus of extremely thermophilic bacteria. Int J Syst Evol Microbiol 1973; 23: 28–36.

10. Stott MB, Crowe MA, Mountain BW, Smirnova A V, Hou S, Alam M, et al. Isolation of novel bacteria, including a candidate division, from geothermal soils in New Zealand. Environ Microbiol 2008; 10: 2030–2041.

11. King CE, King GM. Description of *Thermogemmatispora carboxidivorans* sp. nov., a carbon-monoxide-oxidizing member of the class Ktedonobacteria isolated from a geothermally heated biofilm, and analysis of carbon monoxide oxidation by members of the class Ktedonobacter. Int J Syst Evol Microbiol 2014; 64: 1244–1251.

12. Houghton KM, Morgan XC, Lagutin K, MacKenzie AD, Vyssotski M, Mitchell KA, et al. *Thermorudis pharmacophila* WKT50.2T sp. nov., a novel isolate of class Thermomicrobia isolated from geothermal soil. Int J Syst Evol Microbiol 2015; 65: 4479–4487.

13. King CE, King GM. *Thermomicrobium carboxidum* sp. nov., and *Thermorudis peleae* gen. nov., sp. nov., carbon monoxide-oxidizing bacteria isolated from geothermally heated biofilms. Int J Syst Evol Microbiol 2014; 64: 2586–2592.

14. Spear JR, Walker JJ, McCollom TM, Pace NR. Hydrogen and bioenergetics in the Yellowstone geothermal ecosystem. Proc Natl Acad Sci 2005; 102: 2555–2560.

15. Shock EL, Holland M, Amend JP, Osburn GR, Fischer TP. Quantifying inorganic sources of geochemical energy in hydrothermal ecosystems, Yellowstone National Park, USA. Geochim Cosmochim Acta 2010; 74: 4005–4043.

16. King GM, Weber CF. Interactions between bacterial carbon monoxide and hydrogen consumption and plant development on recent volcanic deposits. ISME J 2008; 2: 195–203.

17. King GM. Contributions of atmospheric CO and hydrogen uptake to microbial dynamics on recent Hawaiian volcanic deposits. Appl Environ Microbiol 2003; 69: 4067–4075.

18. Yang J, Zhou E, Jiang H, Li W, Wu G, Huang L, et al. Distribution and diversity of aerobic carbon monoxide-oxidizing bacteria in geothermal springs of China, the Philippines, and the United States. Geomicrobiol J 2015; 32: 903–913.

19. Wu D, Raymond J, Wu M, Chatterji S, Ren Q, Graham JE, et al. Complete genome sequence of the aerobic CO-oxidizing thermophile Thermomicrobium roseum. PLoS One 2009; 4: e4207.

20. King GM, Weber CF. Distribution, diversity and ecology of aerobic CO-oxidizing bacteria. Nat Rev Microbiol 2007; 5: 107–118.

21. Quiza L, Lalonde I, Guertin C, Constant P. Land-use influences the distribution and activity of high affinity CO-oxidizing bacteria Associated to type I-coxL genotype in soil. Front Microbiol 2014; 5: 271.

22. Greening C, Biswas A, Carere CR, Jackson CJ, Taylor MC, Stott MB, et al. Genomic and metagenomic surveys of hydrogenase diversity indicate H_2_ is a widely-utilised energy source for microbial growth and survival. ISME J 2016; 10: 761–777.

23. Søndergaard D, Pedersen CNS, Greening C. HydDB: a web tool for hydrogenase classification and analysis. Sci Rep 2016; 6: 34212.

24. Ji M, Greening C, Vanwonterghem I, Carere CR, Bay SK, Steen JA, et al. Atmospheric trace gases support primary production in Antarctic desert surface soil. Nature 2017; 552: 400–403.

25. Greening C, Constant P, Hards K, Morales SE, Oakeshott JG, Russell RJ, et al. Atmospheric hydrogen scavenging: from enzymes to ecosystems. Appl Environ Microbiol 2015; 81: 1190–1199.

26. Greening C, Berney M, Hards K, Cook GM, Conrad R. A soil actinobacterium scavenges atmospheric H_2_ using two membrane-associated, oxygen-dependent [NiFe] hydrogenases. Proc Natl Acad Sci U S A 2014; 111.

27. Greening C, Villas-Bôas SG, Robson JR, Berney M, Cook GM. The growth and survival of Mycobacterium smegmatis is enhanced by co-metabolism of atmospheric H_2_. PLoS One 2014; 9: e103034.

28. Berney M, Greening C, Conrad R, Jacobs WR, Cook GM. An obligately aerobic soil bacterium activates fermentative hydrogen production to survive reductive stress during hypoxia. Proc Natl Acad Sci U S A 2014; 111: 11479–11484.

29. Constant P, Chowdhury SP, Pratscher J, Conrad R. Streptomycetes contributing to atmospheric molecular hydrogen soil uptake are widespread and encode a putative high-affinity [NiFe]-hydrogenase. Environ Microbiol 2010; 12: 821–829.

30. Liot Q, Constant P. Breathing air to save energy – new insights into the ecophysiological role of high-affinity [NiFe]-hydrogenase in Streptomyces avermitilis. Microbiologyopen 2016; 5: 47–59.

31. Greening C, Carere CR, Rushton-Green R, Harold LK, Hards K, Taylor MC, et al. Persistence of the dominant soil phylum Acidobacteria by trace gas scavenging. Proc Natl Acad Sci U S A 2015; 112: 10497–10502.

32. Meredith LK, Rao D, Bosak T, Klepac-Ceraj V, Tada KR, Hansel CM, et al. Consumption of atmospheric hydrogen during the life cycle of soil-dwelling actinobacteria. Environ Microbiol Rep 2014; 6: 226–38.

33. Kanno M, Constant P, Tamaki H, Kamagata Y. Detection and isolation of plant-associated bacteria scavenging atmospheric molecular hydrogen. Environ Microbiol 2015; 18: 2495–2506.

34. Khdhiri M, Hesse L, Popa ME, Quiza L, Lalonde I, Meredith LK, et al. Soil carbon content and relative abundance of high affinity H_2_-oxidizing bacteria predict atmospheric H_2_ soil uptake activity better than soil microbial community composition. Soil Biol Biochem 2015; 85: 1–9.

35. Constant P, Poissant L, Villemur R. Isolation of *Streptomyces* sp. PCB7, the first microorganism demonstrating high-affinity uptake of tropospheric H_2_. ISME J 2008; 2: 1066–1076.

36. Myers MR, King GM. Isolation and characterization of *Acidobacterium ailaaui* sp. nov., a novel member of Acidobacteria subdivision 1, from a geothermally heated Hawaiian microbial mat. Int J Syst Evol Microbiol 2016; 66: 5328–5335.

37. Vyssotski M, Ryan J, Lagutin K, Wong H, Morgan X, Stott M. A novel fatty acid, 12,17-dimethyloctadecanoic acid, from the extremophile *Thermogemmatispora* sp. (Strain T81). Lipids 2012; 47: 601–611.

38. Tomazini A, Lal S, Munir R, Stott M, Henrissat B, Polikarpov I, et al. Analysis of carbohydrate-active enzymes in *Thermogemmatispora* sp. strain T81 reveals carbohydrate degradation ability. Can J Microbiol 2018; 10.1139/cjm-2018-0336.

39. Andrews S. FastQC: a quality control tool for high throughput sequence data. 2010.

40. Robinson MD, Oshlack A. A scaling normalization method for differential expression analysis of RNA-seq data. Genome Biol 2010; 11: R25.

41. Rau A, Gallopin M, Celeux G, Jaffrézic F. Data-based filtering for replicated high-throughput transcriptome sequencing experiments. Bioinformatics 2013; 29: 2146–2152.

42. Robinson MD, McCarthy DJ, Smyth GK. edgeR: a Bioconductor package for differential expression analysis of digital gene expression data. Bioinformatics 2010; 26: 139–140.

43. Wentworth WE, Vasnin S V, Stearns SD, Meyer CJ. Pulsed discharge helium ionization detector. Chromatographia 1992; 34: 219–225.

44. Novelli PC, Crotwell AM, Hall BD. Application of gas chromatography with a pulsed discharge helium ionization detector for measurements of molecular hydrogen in the atmosphere. Environ Sci Technol 2009; 43: 2431–2436.

45. Smith PK et al., Krohn R Il, Hermanson GT, Mallia AK, Gartner FH, Provenzano Md, et al. Measurement of protein using bicinchoninic acid. Anal Biochem 1985; 150: 76–85.

46. Walker JM. Nondenaturing polyacrylamide gel electrophoresis of proteins. The protein protocols handbook. 2009. Springer, pp 171–176.

47. Greening C, Berney M, Hards K, Cook GM, Conrad R. A soil actinobacterium scavenges atmospheric H_2_ using two membrane-associated, oxygen-dependent [NiFe] hydrogenases. Proc Natl Acad Sci U S A 2014; 111: 4257–4261.

48. Lorite MJ, Tachil J, Sanjuán J, Meyer O, Bedmar EJ. Carbon monoxide dehydrogenase activity in *Bradyrhizobium japonicum*. Appl Environ Microbiol 2000; 66: 1871–1876.

49. Berney M, Greening C, Hards K, Collins D, Cook GM. Three different [NiFe] hydrogenases confer metabolic flexibility in the obligate aerobe Mycobacterium smegmatis. Environ Microbiol 2014; 16: 318–330.

50. Carere CR, Hards K, Houghton KM, Power JF, McDonald B, Collet C, et al. Mixotrophy drives niche expansion of verrucomicrobial methanotrophs. ISME J 2017; 11: 2599–2610.

51. Altschul SF, Gish W, Miller W, Myers EW, Lipman DJ. Basic local alignment search tool. J Mol Biol 1990; 215: 403–410.

52. Kumar S, Stecher G, Tamura K. MEGA7: Molecular Evolutionary Genetics Analysis version 7.0 for bigger datasets. Mol Biol Evol 2016; msw054.

53. Constant P, Chowdhury SP, Hesse L, Pratscher J, Conrad R. Genome data mining and soil survey for the novel Group 5 [NiFe]-hydrogenase to explore the diversity and ecological importance of presumptive high-affinity H_2_-oxidizing bacteria. Appl Environ Microbiol 2011; 77: 6027–6035.

54. Koch T, Dahl C. A novel bacterial sulfur oxidation pathway provides a new link between the cycles of organic and inorganic sulfur compounds. ISME J 2018; 12: 2479–2491.

55. Cao X, Koch T, Steffens L, Finkensieper J, Zigann R, Cronan JE, et al. Lipoate-binding proteins and specific lipoate-protein ligases in microbial sulfur oxidation reveal an atpyical role for an old cofactor. Elife 2018; 7: e37439.

56. Mohammadi S, Pol A, van Alen TA, Jetten MSM, Op den Camp HJM. Methylacidiphilum fumariolicum SolV, a thermoacidophilic ‘Knallgas’ methanotroph with both an oxygen-sensitive and -insensitive hydrogenase. ISME J 2017; 11: 945–958.

57. Berney M, Cook GM. Unique flexibility in energy metabolism allows mycobacteria to combat starvation and hypoxia. PLoS One 2010; 5: e8614.

58. Cavaletti L, Monciardini P, Bamonte R, Schumann P, Rohde M, Sosio M, et al. New lineage of filamentous, spore-forming, gram-positive bacteria from soil. Appl Environ Microbiol 2006; 72: 4360–4369.

59. Sorokin DY, Vejmelkova D, Lücker S, Streshinskaya GM, Rijpstra WIC, Damste JSS, et al. *Nitrolancea hollandica* gen. nov., sp. nov., a chemolithoautotrophic nitrite-oxidizing bacterium isolated from a bioreactor belonging to the phylum Chloroflexi. Int J Syst Evol Microbiol 2014; 64: 18591865.

60. Klatt CG, Liu Z, Ludwig M, Kühl M, Jensen SI, Bryant DA, et al. Temporal metatranscriptomic patterning in phototrophic Chloroflexi inhabiting a microbial mat in a geothermal spring. ISME J 2013; 7: 1775.

61. Jayachandran G, Görisch H, Adrian L. Studies on hydrogenase activity and chlorobenzene respiration in *Dehalococcoides* sp. strain CBDB1. Arch Microbiol 2004; 182: 498–504.

62. Nijenhuis I, Zinder SH. Characterization of hydrogenase and reductive dehalogenase activities of *Dehalococcoides ethenogenes* strain 195. Appl Environ Microbiol 2005; 71: 1664–1667.

63. Hartwig S, Dragomirova N, Kublik A, Türkowsky D, von Bergen M, Lechner U, et al. A H_2_-oxidizing, 1, 2, 3-trichlorobenzene-reducing multienzyme complex isolated from the obligately organohalide-respiring bacterium *Dehalococcoides mccartyi* strain CBDB1. Environ Microbiol Rep 2017; 9: 618–625.

64. King GM. Molecular and culture-based analyses of aerobic carbon monoxide oxidizer diversity. Appl Environ Microbiol 2003; 69: 7257–65.

65. Weber CF, King GM. The phylogenetic distribution and ecological role of carbon monoxide oxidation in the genus Burkholderia. FEMS Microbiol Ecol 2012; 79: 167–175.

66. King GM. Uptake of carbon monoxide and hydrogen at environmentally relevant concentrations by Mycobacteria. Appl Environ Microbiol 2003; 69: 7266–7272.

67. Ehhalt DH, Rohrer F. The tropospheric cycle of H_2_: a critical review. Tellus B 2009; 61: 500–535.

